# Individual leadership in cattle persists when collective movement is induced by robotic herding

**DOI:** 10.64898/2026.10.14.759049

**Authors:** Hiroki Anzai, Honoka Miyagi, Kanon Tamura, Kirika Iwatani, Sakuya Miyamoto

## Abstract

In social species, some individuals consistently lead collective movements while others follow, but whether these roles persist when movement is externally induced remains unknown. Robotic herding offers a way to address this: animals are displaced by an approaching device, yet retain the option to stay, to move or to follow a conspecific that has moved. We recorded spontaneous group departures in a herd of 18 Japanese Black cows on three days, then recorded which individuals initiated movement away from a quadruped robot, which followed and which remained, across an eight-day herding trial. Cows that frequently initiated spontaneous departures and attracted followers also initiated movement away from the robot more often and drew more followers, whereas habitual followers tended to be less effective initiators. Follower frequency during herding was unrelated to any characteristic measured. Individual leadership therefore persists when movement is induced by a robot. Behavioural roles may thus be expressed under moderate external constraint, provided animals retain some freedom of choice. Identifying leaders from spontaneous departures offers a practical route to more efficient robotic herding.

## 1. Introduction

How collective movement is initiated, and which individuals determine where a group goes, are central questions in the study of collective animal behaviour. Theory shows that a small minority of informed individuals can guide a whole group without explicit signalling, and that the proportion of leaders required decreases as group size increases [1]. Empirically, individuals differ consistently in their tendency to initiate movement or to follow: in white-faced capuchin monkeys, all group members initiated collective movements, and success depended on behavioural cues such as looking back and departing slowly rather than on dominance rank [2]; in chacma baboons, most adults attempted departures, with males initiating more often but without monopoly [3]. Whether leadership is concentrated or distributed, such differences are a repeatable dimension of behavioural variation that shapes how groups move and function [4].

Because spontaneous collective movements cannot be scheduled, it is difficult to observe the same individuals responding repeatedly to a comparable stimulus, or to ask whether their roles remain stable when movement is induced rather than self-generated. A controlled external perturbation offers a solution, and mobile robots are well suited to providing one: they can be deployed repeatedly, directed at specific individuals and made to deliver a consistent stimulus under free-ranging conditions [5,6]. A biomimetic robot accepted by guppies as a conspecific [7] has been used as an experimentally controlled leader to test which traits make an individual followed [8], and a robotic falcon elicited collective escape in free-living bird flocks more reliably than a non-biomimetic drone [9]. Such ethorobotic studies show that robots can serve as controlled, biologically meaningful stimuli in the field, but they have characterized responses at the level of the group, because individuals in wild flocks or shoals are rarely identifiable. Whether individuals occupy the same roles in undisturbed and robot-induced movement therefore remains untested; answering this requires identified individuals observed under both conditions.

Whether roles persist under induced movement is not a foregone conclusion. In cattle and sheep driven by human handlers, the order of animals during forced movement was unrelated to that during voluntary movement [10–12]. In those studies, however, animals were routed along fences, driven into dipping tanks or moved through handling facilities, where spatial constraint and human pressure left almost no room for individual choice; movement order may therefore have reflected physical position and crowding rather than behavioural disposition. Robotic herding in an open pasture is intermediate on this continuum of behavioural freedom: animals are displaced by an approaching device, yet can still stay, move or follow a conspecific that has moved. Whether leadership and followership survive this intermediate level of constraint is therefore an open question, and one that bears on the conditions under which behavioural roles are expressed at all. A second question is whether leadership is consistent across motivationally distinct contexts. Under the ‘leading according to need’ framework, the individual with the greatest immediate requirement initiates movement, so leaders should change as the motivation for moving changes [13]; if leadership instead reflects stable individual attributes, the same individuals should lead irrespective of context [14,15].

Cattle offer a system in which both questions can be addressed. Herds are of manageable size, individuals can be marked and followed for months, and spontaneous group departures are frequent enough to characterize each individual’s tendency to initiate or to follow before a robot is introduced. Individuals that consistently occupy forward positions in a moving herd and whose departures attract followers are recognized as leaders, and those that consistently follow as followers [16,17]. These roles matter at the group level: replacing a follower cow with a leader changes how a herd distributes itself across a pasture [18]. Leadership in cattle is associated with high dominance rank [19], and dominant cattle spend less time grazing [20,21]. In baboons, leadership is also embedded in social relationships: individuals central to the network of spatial associations are more often followed, irrespective of their dominance rank [22].

Grazing animals use pasture selectively, causing soil degradation, loss of plant species diversity and inefficient forage utilization [23]. Robotic herding, the use of autonomous or semi-autonomous devices to direct animal movement, has attracted attention as a labour-saving way to make grazing distribution more uniform [5,24]: ground-based unmanned vehicles can steer cattle away from intensively grazed patches without inducing strong aversion [25], whereas cattle respond weakly to drones and habituate rapidly to them [26]. In a companion study of the same herd [27], a quadruped robot elicited stronger responses and higher herding success than a drone. Herding nevertheless remained effective on some days when individual responsiveness had declined, possibly because responsive individuals that moved away drew along conspecifics that had become less reactive. Because following was not analysed at the individual level, this explanation could not be tested.

Here we used a quadruped robot as a controlled perturbation to ask whether the leadership and followership roles that individuals express during spontaneous collective movement are also expressed when movement is externally induced. Within the same experimental series, we characterized 18 Japanese Black cows by their behaviour during spontaneous group departures and their feeding position within the herd, and then recorded which individuals initiated movement away from the robot, and which followed or stayed, over an eight-day herding trial. We predicted that individuals with stronger leadership during spontaneous movement would also initiate movement away from the robot and attract more followers, whereas individuals with stronger followership would be less effective initiators and more likely to follow others during herding. We also examined whether time spent near the group but not grazing, a behavioural correlate of dominance [20,21], feeding position and age were related to the roles individuals adopted during herding.

## 2. Material and methods

### 2.1. Study site and animals

This study was conducted as part of the same experimental series described in the companion study [27], to which the reader is referred for full details of the study site and animal management. Briefly, trials were conducted from May to October 2023 at the Sumiyoshi Livestock Science Station (31°59′N, 131°28′E), Faculty of Agriculture, University of Miyazaki. The 1.1 ha experimental pasture was divided into nine equal plots in a 3 × 3 grid (A1–C3; see [27], figure 1 for layout). A herd of 25–29 Japanese Black breeding cows and 7– 11 calves was stocked on the experimental pasture for five consecutive days each month (09:00–16:00 h). To enable individual-level behavioural tracking, numbers were painted on the body of each adult cow using livestock marker paint (Tell Tail Aerosols, FIL, New Zealand) before each grazing period. All procedures were approved by the Animal Care and Use Committee of the University of Miyazaki (approval number 2020-006-5).

**Figure 1.**
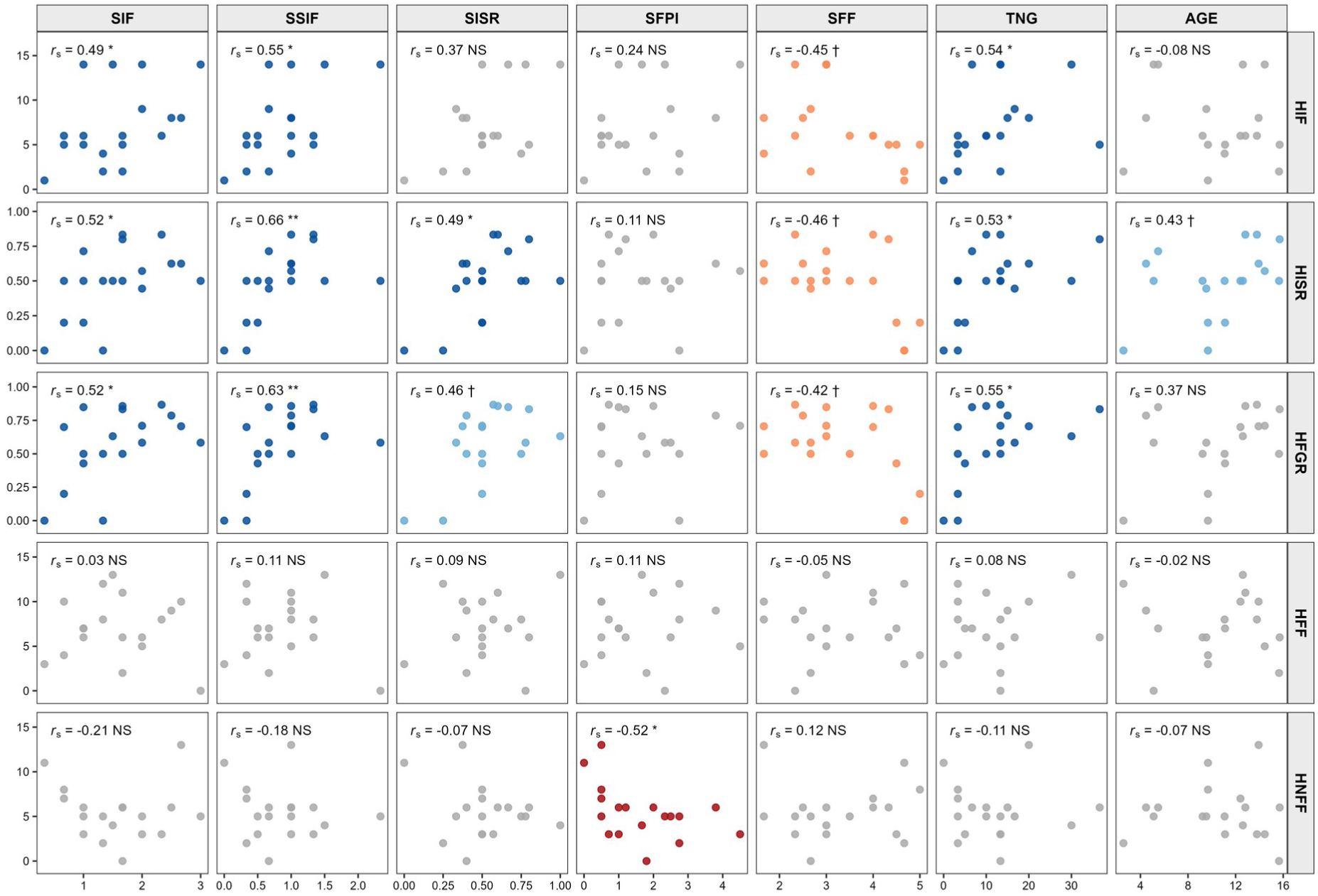
Scatter plot matrix showing relationships between individual characteristics during spontaneous collective movement (columns: SIF, initiator frequency; SSIF, initiation success frequency; SISR, initiation success rate; SFPI, average followers per initiation; SFF, follower frequency; TNG, time spent near the group but not grazing; AGE, age) and individual responses during robotic herding (rows: HIF, initiator frequency; HISR, initiation success rate; HFGR, follower generation rate; HFF, follower frequency; HNFF, non-follower frequency). Spearman’s rank correlation coefficient (*r*_s_) and significance level are shown in the upper-left corner of each panel. NS, not significant; †, *P* < 0.1; *, *P* < 0.05; **, *P* < 0.01; ***, *P* < 0.001. *n* = 18.

### 2.2. Individual characteristics during spontaneous collective movement

To characterize individual tendencies in spontaneous collective movement, behavioural observations were conducted on the first grazing day of each month in May, June and July by two observers working simultaneously.

One observer recorded voluntary group departures, defined as occasions when cattle raised their heads from grazing and moved towards the resting area or shade trees. For each departure, the observer recorded the identity of the first individual to initiate movement (initiator) and the identities of individuals that followed the initiator within 1 min (followers). An initiation was recorded as successful if at least one follower joined the initiator, and as unsuccessful if no individual followed within 1 min.

The second observer recorded the feeding positions of all individuals at 10 min intervals throughout the grazing period. At each interval, individuals were ordered from front to rear along the predominant direction of group movement, defined as the direction faced by the majority of individuals. Intervals during which more than one-third of individuals were not grazing, or no clear group direction was discernible, were excluded. At each 10 min scan, individuals located within five cow-lengths of the nearest group member but not grazing were also recorded; the number of such scans was converted to minutes per day by multiplying by the 10 min scan interval.

Based on the 10 min interval data, each individual was assigned a feeding position type over the three observation days. Individuals were classified as front cows if their mean daily order was consistently within the top 10 on at least two of the three days. Individuals were classified as rear cows if their mean daily rank was consistently within the bottom 10 on at least two of the three days. Individuals were classified as middle cows if they were consistently ranked within four to five positions of the median rank on at least two of the three days.

The variables derived from these observations are listed in table 1.

**Table 1.**
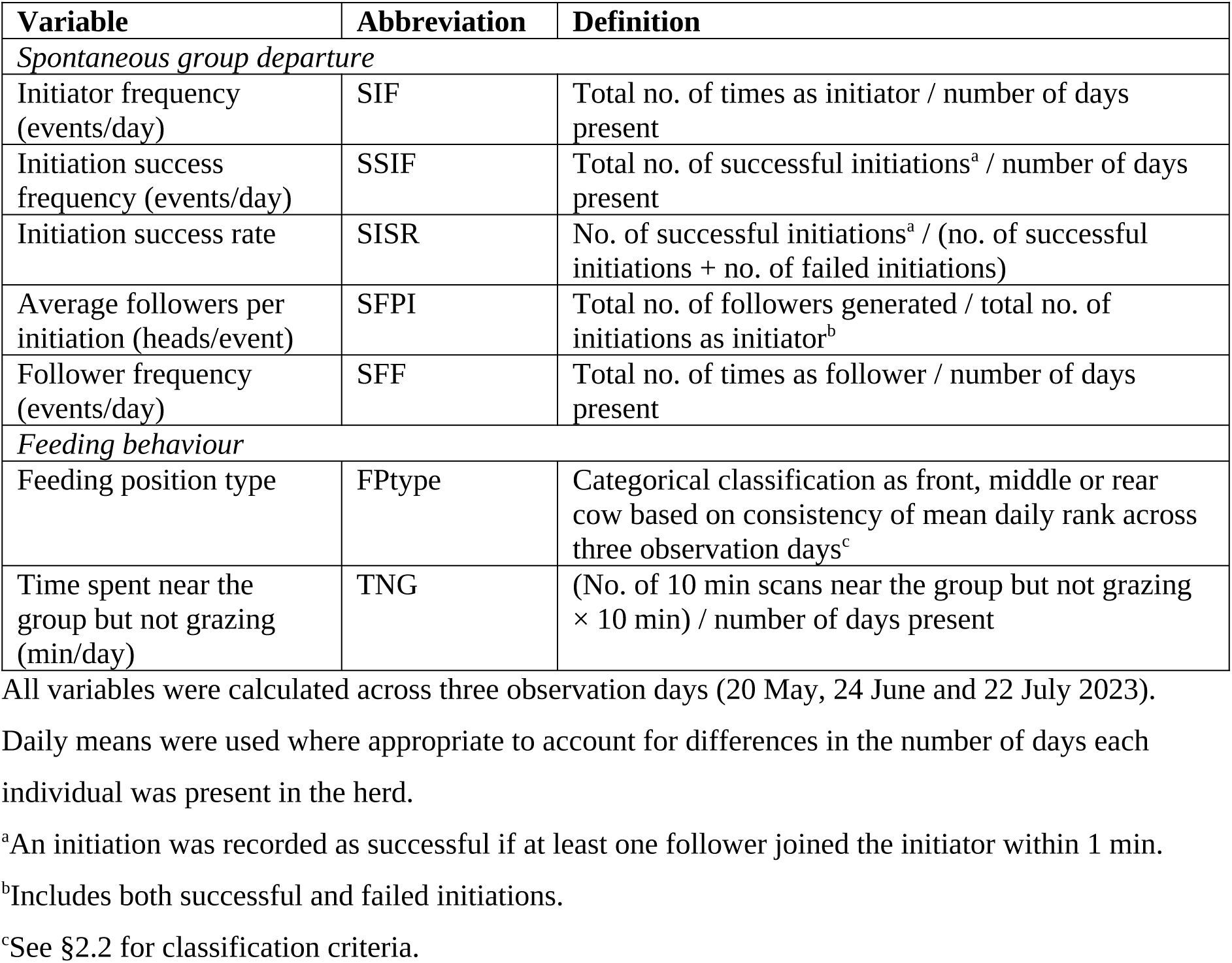
Variables describing individual characteristics during spontaneous collective movement.

| Variable | Abbreviation | Definition |
| --- | --- | --- |
| *Spontaneous group departure* |  |  |
| Initiator frequency (events/day) | SIF | Total no. of times as initiator / number of days present |
| Initiation success frequency (events/day) | SSIF | Total no. of successful initiations<sup>a</sup> / number of days present |
| Initiation success rate | SISR | No. of successful initiations<sup>a</sup> / (no. of successful initiations + no. of failed initiations) |
| Average followers per initiation (heads/event) | SFPI | Total no. of followers generated / total no. of initiations as initiator<sup>b</sup> |
| Follower frequency (events/day) | SFF | Total no. of times as follower / number of days present |
| *Feeding behaviour* |  |  |
| Feeding position type | FPtype | Categorical classification as front, middle or rear cow based on consistency of mean daily rank across three observation days<sup>c</sup> |
| Time spent near the group but not grazing (min/day) | TNG | (No. of 10 min scans near the group but not grazing × 10 min) / number of days present |

All variables were calculated across three observation days (20 May, 24 June and 22 July 2023).
Daily means were used where appropriate to account for differences in the number of days each individual was present in the herd.
<sup>a</sup>An initiation was recorded as successful if at least one follower joined the initiator within 1 min.
<sup>b</sup>Includes both successful and failed initiations.
<sup>c</sup>See §2.2 for classification criteria.

### 2.3. Robotic herding trials and individual response measurements

Robotic herding trials were conducted on all five grazing days in September and the first three grazing days in October (eight days in total), following the procedure described in Anzai *et al.* [27]. A quadruped robot (Unitree Go1; Unitree Robotics Co., Ltd., Zhejiang, China) equipped with a speaker delivering dog-barking sounds was operated by a single operator throughout the trials. The herding area comprised three adjacent plots in a single column along one edge of the pasture (A1, B1 and C1; approximately 33% of the pasture area). A herding session was initiated whenever two or more cows were observed grazing within the herding area. The operator manoeuvred the robot to approach a target cow from outside its rear blind angle; if the cow did not move away when the robot came within approximately one cow-length, the robot performed lateral movements, rotations and vertical head motions for approximately 20 s. If the target cow remained unresponsive, the operator switched to another cow grazing within the herding area. A session ended when all cows had left the herding area or when all targeted cows remained unresponsive for 15–20 min. Calves were not targeted. All sessions were video-recorded (HDR-CX180; Sony, Tokyo, Japan).

From the video records, the following were identified for each herding event: the first individual to respond to the approaching robot and move out of the herding area (initiator); individuals that left the herding area following the initiator’s departure (followers); and individuals that remained in the herding area despite being within one cow-length of the initiator at the time of its departure (non-followers). An initiation was recorded as successful if at least one follower was observed, and as unsuccessful if no individual followed.

The variables derived from these records are listed in table 2.

**Table 2.**
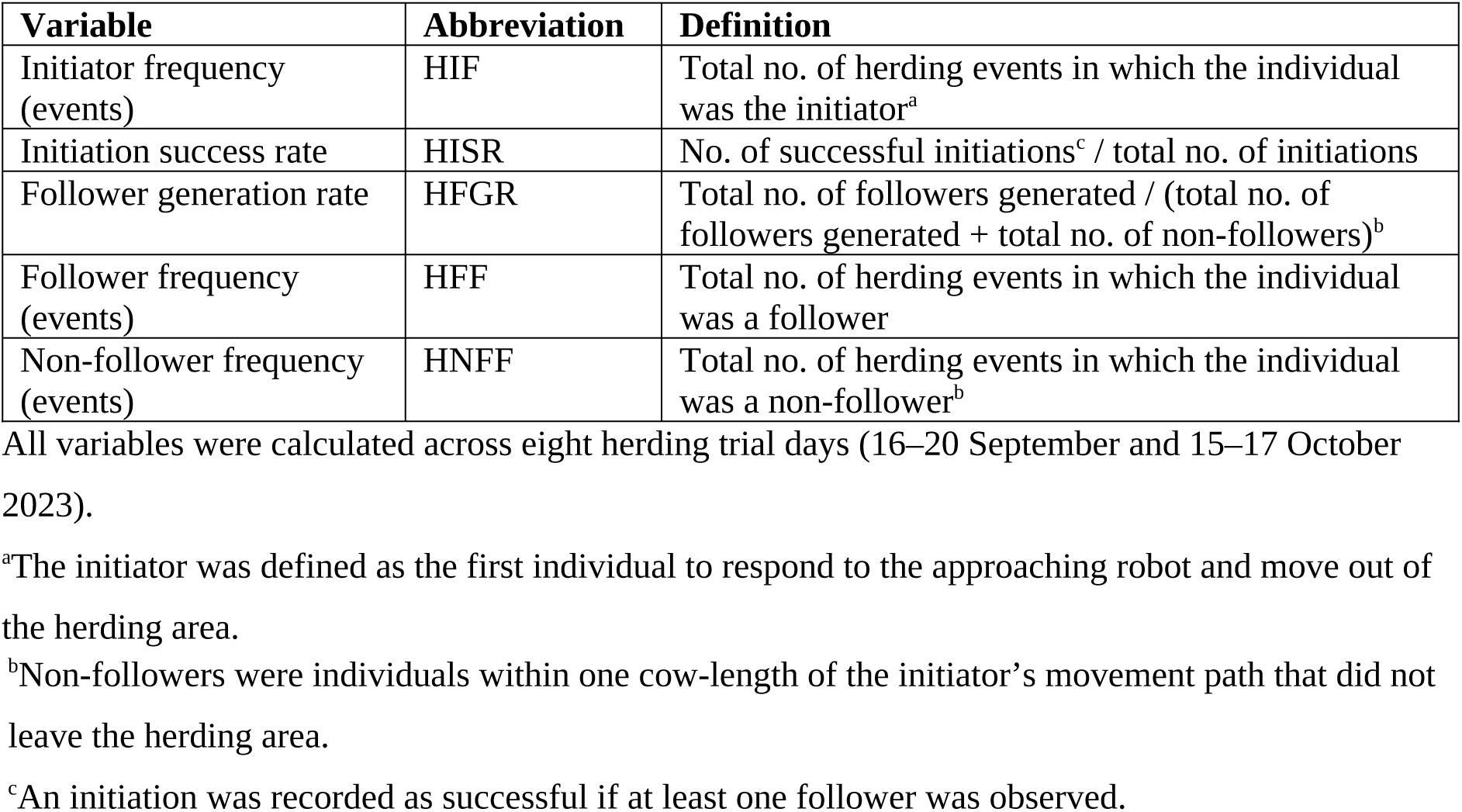
Variables describing individual behaviour during robotic herding.

| Variable | Abbreviation | Definition |
| --- | --- | --- |
| Initiator frequency (events) | HIF | Total no. of herding events in which the individual was the initiator<sup>a</sup> |
| Initiation success rate | HISR | No. of successful initiations<sup>c</sup> / total no. of initiations |
| Follower generation rate | HFGR | Total no. of followers generated / (total no. of followers generated + total no. of non-followers)<sup>b</sup> |
| Follower frequency (events) | HFF | Total no. of herding events in which the individual was a follower |
| Non-follower frequency (events) | HNFF | Total no. of herding events in which the individual was a non-follower<sup>b</sup> |

All variables were calculated across eight herding trial days (16–20 September and 15–17 October 2023).
<sup>a</sup>The initiator was defined as the first individual to respond to the approaching robot and move out of the herding area.
<sup>b</sup>Non-followers were individuals within one cow-length of the initiator's movement path that did not leave the herding area.
<sup>c</sup>An initiation was recorded as successful if at least one follower was observed.

### 2.4. Statistical analysis

Of the cows present during the study, only individuals present in the herd for at least two of the three monthly observation periods in May–July and for all eight herding trial days in September and October were included in the analysis (*n* = 18). For variables related to spontaneous collective movement, daily means were used where appropriate to account for differences in the number of observation days among individuals.

Relationships among variables related to spontaneous collective movement and robotic herding were evaluated using Spearman’s rank correlation coefficient (*r*_s_), as normality of the data was not assumed given the small sample size (*n* = 18). The age of each individual was also examined for correlations with all other quantitative variables, to evaluate whether age was associated with individual movement roles during either spontaneous collective movement or robotic herding. Differences in herding-related variables among feeding position types (front, middle and rear cows) were assessed using the Kruskal–Wallis test followed by the Steel–Dwass multiple comparison test. The two individuals that could not be assigned to any feeding position type were excluded from this analysis but were included in the correlation analyses. All analyses were conducted using SAS version 9.4 (SAS Institute Inc., Cary, NC, USA; PROC NPAR1WAY and PROC CORR). The significance level was set at 0.05 for all tests. Because the present analysis was exploratory in nature, correlations were evaluated without correction for multiple testing, and the results should therefore be regarded as indicating patterns to be confirmed in future work. A scatter plot matrix and box plots were produced using R version 4.5.3 [28], with the GGally [29] and ggplot2 [30] packages.

## 3. Results

Descriptive statistics for all variables are shown in table 3. Several variables related to spontaneous collective movement were correlated with initiator-related variables during robotic herding (figure 1). Initiator frequency (SIF) and initiation success frequency (SSIF) during spontaneous departures were positively correlated with initiator frequency (HIF), initiation success rate (HISR) and follower generation rate (HFGR) during herding (*r*_s_ = 0.49 to 0.66, *P* < 0.05). Follower frequency (SFF) showed negative trends with HIF, HISR and HFGR (*r*_s_ = −0.46 to −0.42, *P* < 0.1). Time spent near the group but not grazing (TNG) was positively correlated with HIF, HISR and HFGR (*r*_s_ = 0.53 to 0.55, *P* < 0.05). In contrast, follower frequency during herding (HFF) was not correlated with any individual characteristic examined (*P* > 0.1). Non-follower frequency during herding (HNFF) was negatively correlated only with the average followers per initiation during spontaneous departures (SFPI; *r*_s_ = −0.52, *P* < 0.05). Age was not significantly correlated with any herding response variable, except for a positive trend with HISR (*r*_s_ = 0.43, *P* < 0.1), nor with any variable describing spontaneous collective movement (*r*_s_ = −0.14 to 0.39, *P* > 0.1).

**Table 3.** Descriptive statistics of age and variables related to collective movement and robotic herding (*n* = 18).

| Variable | Median | (min–max) |
| --- | --- | --- |
| *Variables related to collective movement* |  |  |
| Initiator frequency (events/day) | 1.583 | (0.333–3.000) |
| Initiation success frequency (events/day) | 0.833 | (0.000–2.333) |
| Initiation success rate | 0.500 | (0.000–1.000) |
| Average followers per initiation (heads/event) | 1.433 | (0.000–4.500) |
| Follower frequency (events/day) | 3.000 | (1.667–5.000) |
| Time spent near the group but not grazing (min/day) | 11.67 | (0.00–36.67) |
| *Variables related to robotic herding* |  |  |
| Initiator frequency (events) | 6.0 | (1.0–14.0) |
| Initiation success rate | 0.500 | (0.000–0.833) |
| Follower generation rate | 0.607 | (0.000–0.867) |
| Follower frequency (events) | 7.0 | (0.0–13.0) |
| Non-follower frequency (events) | 5.0 | (0.0–13.0) |
| *Individual attribute* |  |  |
| Age (years) | 11.09 | (2.57–15.70) |

Of the 18 cows included in the analysis, 5 were classified as front cows, 5 as middle cows and 6 as rear cows; the remaining 2 could not be assigned to any category. Initiator frequency during robotic herding differed significantly among feeding position types (Kruskal–Wallis: *χ*^2^ = 6.88, d.f. = 2, *P* = 0.032), and was higher in middle cows than in rear cows (Steel– Dwass: *P* = 0.034; figure 2). No significant differences among feeding position types were detected for the remaining herding response variables.

**Figure 2.**
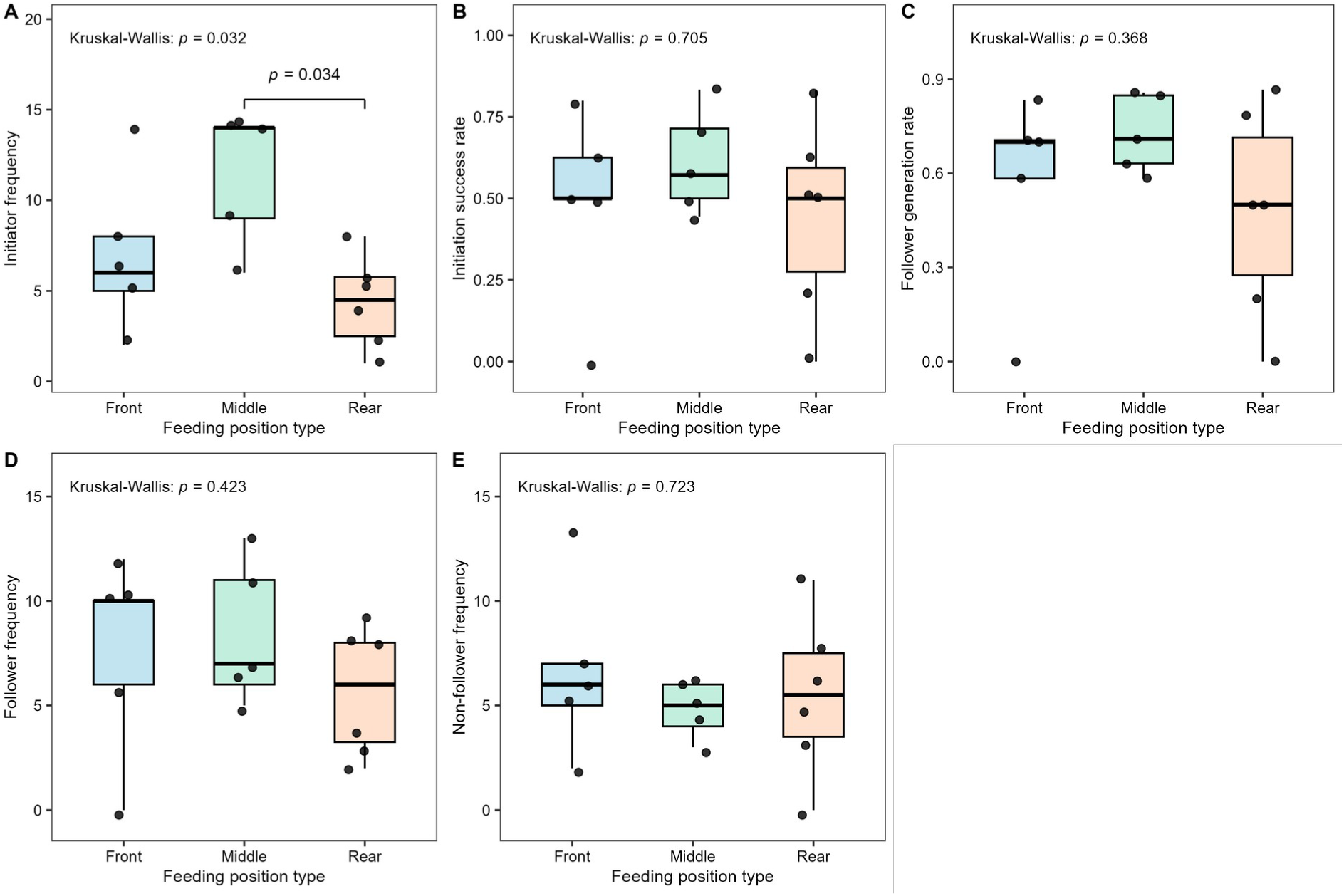
Individual responses during robotic herding by feeding position type (front, *n* = 5; middle, *n* = 5; rear, *n* = 6). (A) Initiator frequency; (B) initiation success rate; (C) follower generation rate; (D) follower frequency; (E) non-follower frequency. Kruskal–Wallis *P*-values are shown above each panel. Brackets indicate pairwise differences assessed by the Steel–Dwass test.

## 4. Discussion

Cows that frequently initiated spontaneous departures and attracted followers, that is, those with strong leadership [16,17], also initiated movement away from the robot more often and drew conspecifics along. This supports the suggestion of the companion study [27] that individual variation in leadership contributed to the unexpectedly high herding success on the third day of continuous trials.

Conversely, cows that frequently followed others during spontaneous departures tended to be less effective initiators during herding, with little influence on the movements of nearby conspecifics. Similarly, a herd containing a cow with strong followership exploited a more restricted area of pasture than the same herd in which that cow was replaced by one with strong leadership [18], indicating that followers do not draw conspecifics into new areas.

These roles persisted even though movement was induced, whereas studies using human handlers found no relationship between voluntary and forced movement orders [10–12]. Those forced movements involved fences, dipping tanks or handling facilities, which severely constrained individual choice. In the present study, cattle could remain in the herding area and keep grazing even when approached by the robot, making herding a semi-forced movement, intermediate on the continuum of behavioural freedom described in the Introduction. This freedom may have been a prerequisite for leadership and followership tendencies to influence herding outcomes.

The two contexts also differed in motivation: spontaneous departures were associated with foraging and resting, whereas movement during herding was elicited by an aversive stimulus. Under the ‘leading according to need’ framework, leadership should therefore have shifted between contexts [13]. That the same individuals led in both suggests instead that leadership in this herd rested on stable attributes rather than transient state. This accords with the view that leadership is assumed consistently by particular individuals in groups with stable membership and established dominance hierarchies [14], and with reports that consistent individual differences predict leadership across settings [15]. The aversive context examined here was mild, however: overt fear responses to the robot were rare [27], and whether leadership is conserved under a stronger threat remains to be tested. Consistent leadership also need not reflect a dedicated tendency to lead. Where positions and following arise from social relationships, consistent roles can emerge as by-products, or ‘social spandrels’ [31,32], and may then carry over to a novel, human-imposed context such as robotic herding.

Two of the characteristics measured here offer clues to what these stable attributes might be. The first is time spent near the group but not grazing, which was positively associated with all three initiator-related variables during herding. This measure may reflect dominance: dominant cattle graze less and ruminate more than subordinates [20,21], presumably because priority of access to preferred patches lets them meet their intake requirements more efficiently, and an individual’s influence on group movement increases with its dominance rank [19]. Alternatively, the measure may index social centrality: in chacma baboons, individuals central to the network of spatial associations were more often followed at group departures, whereas dominance rank did not predict initiation success [22]. Either way, cows with high values may have been individuals whose departures nearby animals readily attended to, although neither dominance rank nor social relationships were measured here.

The second is position within the grazing herd. Contrary to our expectation that front cows would initiate most often, middle cows initiated more often than rear cows, so feeding position does not appear to reflect leadership directly. Leadership in cattle is best measured during long-distance spontaneous movements towards a new feeding site, and movement order during grazing bouts follows a different distribution [16]. Position during grazing may instead reflect a tendency to stay near conspecifics: middle cows consistently occupied central positions within the grazing group and, when displaced by the robot, may have set nearby conspecifics in motion. Similarly, dominant baboons occupy central positions during travel because they have more close associates, not because they lead [31], and well-connected, centrally positioned individuals are more often followed [22]. Together with the result for time spent near the group, this suggests that social centrality may underlie leadership in both contexts. This could be tested by measuring social relationships and spatial positions directly: if differences in leadership disappeared after controlling for these factors, leadership would reflect social centrality rather than an individual tendency to lead.

Age, by contrast, was not significantly correlated with any herding variable or with leadership during spontaneous movement, suggesting that behavioural tendencies rather than age determined movement roles during herding. In a semi-wild cattle herd, travelling position during voluntary movement was likewise independent of age, with an age-related order emerging only during forced movement [11], and age did not predict initiation success in baboons [22]. This is notable given the wide age range of the present herd (2.6–15.7 years).

Following showed a different pattern. Follower frequency during herding was unrelated to any characteristic measured, suggesting that the tendency to follow a displaced conspecific does not depend on an individual’s role in spontaneous movement. In cattle, initiation and following are not restricted to particular individuals [33], and in baboons and goats, group members follow the movement of initiators rather than dedicated signals [22,34]; following may therefore require little individual assessment of the stimulus. Non-follower frequency, by contrast, was lower in cows that attracted more followers during spontaneous departures. Individuals linked by social bonds frequently follow each other in cattle and sheep [35], and cattle synchronize their activities, particularly on pasture [33]. Strong leaders, being socially central, may therefore be drawn into collective movement when others leave, whereas less integrated individuals more often remain behind.

Methodologically, robotic perturbation can make the study of collective movement more efficient. Spontaneous departures occur only when animals decide to move, so observers must wait for them, whereas a robot can induce movement on demand. Because the roles individuals adopted during induced movement were associated with those during spontaneous departures, robot-induced movement may help identify leaders and followers without waiting for spontaneous events. This illustrates the dual value of ethorobotics noted by Storms *et al.* [9]: the same robot serves both as a controlled stimulus for studying collective movement and as a tool for managing where animals go. In practice, targeting strong leaders could improve herding efficiency by moving several animals at once, a strategy also proposed for free-roaming ungulate herds [36]. Whereas robotic predators have been developed to deter bird flocks, for example around airfields [37], and to induce costly antipredator responses in invasive fish [38], a quadruped robot used with livestock aims not to remove animals from an area but to redistribute them within it, and knowing which individuals carry the group with them makes such fine-grained control conceivable. Leaders could be identified from spontaneous departures or, more simply, from time spent near the group but not grazing.

This study has several limitations. First, it was based on 18 cows from a single herd in a small pasture, and whether the same associations hold in larger herds, other breeds or other pasture conditions remains to be investigated. The small sample also limits statistical power, so the absence of associations for follower frequency during herding should not be taken as evidence that none exist. Second, movement roles were assessed on only three days (May– July), and their stability across seasons and herding contexts was not evaluated. Third, whether leaders can be identified reliably from shorter observation periods, as farm management would require, warrants further investigation. Finally, the causal direction of the association between time spent near the group but not grazing and the initiator-related variables could not be determined. Dominant, socially central or leading individuals may have stayed near the group for reasons unrelated to herding, or their proximity may have resulted from others following them. Moreover, because such individuals were often surrounded by conspecifics, their own displacement by the robot may have been more likely to set others in motion. Separating these possibilities would require tracking the fine-scale movements of individuals, which was beyond the scope of this observational study.

## 5. Conclusion

Cattle that led spontaneous group departures also led when movement was induced by a quadruped robot, initiating movement away from it and drawing conspecifics along, which supports the explanation proposed in the companion study [27]. Initiation during herding was associated with time spent near the group but not grazing and with a central feeding position, but not with age, consistent with leadership grounded in social dominance or social centrality. Following, by contrast, was largely independent of individual characteristics, except that cows that attracted many followers were less likely to remain behind when others left. Individual roles in collective movement therefore persist under moderate external constraint. Because a robot can induce collective movement on demand, robotic perturbation offers an efficient way to study these roles. In practice, leaders could be identified from spontaneous departures or, more simply, from time spent near the group but not grazing, and targeting them may make robotic herding more efficient.

## Ethics

All procedures were approved by the Animal Care and Use Committee of the University of Miyazaki (approval number 2020-006-5) and were carried out in accordance with the ARRIVE guidelines and relevant institutional and national regulations on the care and use of animals. At the end of the study, all animals were returned to the normal management of the research station; no animals were euthanized or harmed as part of the experiment.

## Data accessibility

The data supporting this article are openly available from Zenodo [39] (https://doi.org/10.5281/zenodo.23357826). Behavioural response data from the same experimental series are available from Zenodo [40] (https://doi.org/10.5281/zenodo.20825305). The SAS code used for all analyses is provided in the electronic supplementary material.

## Declaration of AI use

During the preparation of this manuscript, the authors used Claude (Anthropic) to assist with English-language editing, with reorganizing the framing of the Introduction and Discussion and with writing the R code used to generate the figures. The tool was not used to generate scientific insights, to analyse or interpret the data or to draw conclusions; all statistical analyses were performed by the authors, and all figures produced with the assisted code were checked by the authors against the underlying data. After using this tool, the authors reviewed and edited the content as needed and take full responsibility for the content of the published article.

## Authors’ contributions

H.A.: conceptualization, methodology, formal analysis, investigation, data curation, writing —original draft, writing—review and editing, visualization, supervision, project administration, funding acquisition; H.M.: conceptualization, investigation, data curation, writing—original draft; K.T.: investigation, data curation, writing—original draft; K.I.: investigation, data curation, writing—original draft; S.M.: investigation, data curation, writing—original draft.

All authors gave final approval for publication and agreed to be held accountable for the work performed therein.

## Acknowledgements

We thank Ikuo Kobayashi, Genki Ishigaki, Koichiro Henmi and the staff of the Sumiyoshi Livestock Science Station, University of Miyazaki (Miyazaki, Japan), where this study was conducted, for animal management and field assistance, and Aya Nishiwaki for lending us the drone. We also thank Yosuke Goto, Minori Kan, Mizuki Tagashira, Moe Matsuhashi and Yui Kawano for field support.

